# The germline mutational process in rhesus macaque and its implications for phylogenetic dating

**DOI:** 10.1101/2020.06.22.164178

**Authors:** Lucie A. Bergeron, Søren Besenbacher, Jaco Bakker, Jiao Zheng, Panyi Li, George Pacheco, Mikkel-Holger S. Sinding, Maria Kamilari, M. Thomas P. Gilbert, Mikkel H. Schierup, Guojie Zhang

## Abstract

Understanding the rate and pattern of germline mutations is of fundamental importance for understanding evolutionary processes. Here we analyzed 19 parent-offspring trios of rhesus macaques (*Macaca mulatta*) at high sequencing coverage of ca. 76X per individual, and estimated an average rate of 0.77 × 10^−8^ *de novo* mutations per site per generation (95 % CI: 0.69 × 10^−8^ - 0.85 × 10^−8^). By phasing 50 % of the mutations to parental origins, we found that the mutation rate is positively correlated with the paternal age. The paternal lineage contributed an average of 81 % of the *de novo* mutations, with a trend of an increasing male contribution for older fathers. About 3.5 % of *de novo* mutations were shared between siblings, with no parental bias, suggesting that they arose from early development (postzygotic) stages. Finally, the divergence times between closely related primates calculated based on the yearly mutation rate of rhesus macaque generally reconcile with divergence estimated with molecular clock methods, except for the Cercopithecidae/Hominoidea molecular divergence dated at 52 Mya using our new estimate of the yearly mutation rate.

## Introduction

Germline mutations are the source of heritable disease and evolutionary adaptation. Thus, having precise estimates of germline mutation rates is of fundamental importance for many fields in biology, including searching for *de novo* disease mutations (Acuna-Hidalgo et al. 2016; Oliveira et al. 2018), inferring demographic events (Lapierre et al. 2017; Zeng et al. 2018), and accurate dating of species divergence times (Teeling et al. 2005; Ho and Larson 2006; Pulquério and Nichols 2007). Over the past ten years, new sequencing techniques have allowed deep sequencing of individuals from the same pedigree, enabling direct estimation of the *de novo* mutation rate for each generation, and precise estimation of the individual parental contributions to germline mutations across the whole genome. Most such studies have been conducted on humans, using large pedigrees with up to 3000 trios (Jónsson et al. 2017; Halldorsson et al. 2019), leading to a consensus estimate of ∽1.25 x 10^−8^ *de novo* mutation per site per generation, with an average parental age of ∽ 29 years, leading to a yearly rate of 0.43 x 10^−9^ *de novo* mutation per site per year and most variation between trios explained by the age of the parents (Awadalla et al. 2010; Roach et al. 2010; Kong et al. 2012; Neale et al. 2012; Wang and Zhu 2014; Besenbacher et al. 2015; Rahbari et al. 2016; Jónsson et al. 2017; Maretty et al. 2017). The observed increases in the mutation rate with paternal age in humans and other primates (Venn et al. 2014; Jónsson et al. 2017; Thomas et al. 2018) has generally been attributed to errors during replication (Li et al. 1996; Crow 2000). In mammalian spermatogenesis, primordial germ cells go through meiotic divisions, to produce stem cells by the time of puberty. After this time, stem cell divisions occur continuously throughout the male lifetime. Thus, human spermatogonial stem cells have undergone 100 to 150 mitoses in a 20 years old male, and ∽ 610 mitoses in a 40 years old male (Acuna-Hidalgo et al. 2016), leading to an additional 1.51 *de novo* mutations per year increase in the father’s age (Jónsson et al. 2017). Female age also seems to affect the mutation rate in humans, with 0.37 mutations added per year (Jónsson et al. 2017). This maternal effect cannot be attributed to replication errors, as different from spermatogenesis, female oocytogenesis occurs during embryogenesis process and is already finished before birth (Byskov 1986). Moreover, there seems to be a bias towards males in contribution to *de novo* mutations, as the paternal to maternal contribution is 4:1 in human and chimpanzee (Venn et al. 2014; Jónsson et al. 2017). One recent study proposed that damage-induced mutations might be a potential explanation for the observation of both the maternal age effect and the male-bias also present in parents reproducing right after puberty when replication mutations should not have accumulated yet in the male germline (Gao et al. 2019). Parent-offspring analyses can also be used to distinguish mutations that are caused by gametogenesis from mutations that emerge in postzygotic stages (Acuna-Hidalgo et al. 2015; Scally 2016). While germline mutations in humans are relatively well studied, it remains unknown how much variability exists among primates on the contribution of replication errors to *de novo* mutations, the parental effects, and the developmental stages at which these mutations are established (postzygotic or gametogenesis).

Up until now, the germline mutation rate has only been estimated using pedigrees in few non-human primate species, including chimpanzee (*Pan troglodytes*) (Venn et al. 2014; Tatsumoto et al. 2017; Besenbacher et al. 2019), gorilla (*Gorilla gorilla*) (Besenbacher et al. 2019), orangutan (*Pongo abelii*) (Besenbacher et al. 2019), African green monkey (*Chlorocebus sabaeus*) (Pfeifer 2017), owl monkey (*Aotus nancymaae*) (Thomas et al. 2018) and recently rhesus macaque (*Macaca mulatta*) (Wang et al. 2020). The mutation rate of baboon (*Papio anubis*) (Wu et al. 2019) and grey mouse lemur (*Microcebus murinus*) (Campbell et al. 2019) have also been estimated in preprinted studies. To precisely call *de novo* mutations in the offspring, collecting and comparing the genomic information of the pedigrees is a first essential step for detecting mutations only present in offspring but not in either parent. Next, the *de novo* mutations need to be separated from sequencing errors or somatic mutations, which cause false-positive calls. Because mutations are rare events, detecting *de novo* mutations that occur within a single generation requires high sequencing coverage in order to cover a majority of genomic regions and identify the false-positives. Furthermore, the algorithms used to estimate the mutation rate should take false-negative calls into account. However, a considerable range of sequencing depth (ranging from 18X (Pfeifer 2017) to 120X (Tatsumoto et al. 2017)) has been applied in many studies for estimation of mutation rate. Different filtering methods have been introduced to reduce false-positives and false-negatives but the lack of standardized methodology makes it difficult to assess whether differences in mutation rate estimates are caused by technical or biological variability. In addition, most studies on non-human primates used small pedigrees with less than ten trios, which made it difficult to detect any statistically significant patterns over *de novo* mutation spectra.

Studying non-human primates could help us understanding whether the mutation rate is affected by life-history traits such as mating strategies or the age of reproduction. The variation in mutation rate among primates will also be useful for re-calibrating the speciation times across lineages. The sister group of Hominoidea is Cercopithecidae, including the important biomedical model species, rhesus macaque (*Macaca mulatta*), which share 93 % of its genome with humans (Gibbs et al. 2007). This species has a generation time estimate of ∽ 11 years (Xue et al. 2016), and their sexual maturity is much earlier than in humans with females reaching maturity around three years old, while males mature around the age of 4 years (Rawlins and Kessler 1986). While female macaques generally start reproducing right after maturation, males rarely reproduce in the wild until they reach their adult body size, at approximately eight years old (Bercovitch et al. 2003). They are also a promiscuous species, and do not form pair bonds, but reproduce with multiple individuals. These life-history traits, along with their status as the closest related outgroup species of the hominoid group, make the rhesus macaque an interesting species for investigating the differences and common features in mutation rate processes across primates. In this study, we, produced high depth sequencing data for 33 rhesus macaque individuals (76X per individual) representing 19 trios. This particular dataset consists of a large number of trios, each with high coverage sequencing, and allowed us to test different filter criteria and choose the most appropriate ones to estimate the species mutation rate with high confidence. With a large number of *de novo* mutations phased to their parents of origins, we can statistically assess the parental contribution and the effect of the parental age. We characterize the type of mutations and their location on the genome to detect clusters and shared mutations between siblings. Finally, we use our new estimate to infer the effective population size and date their divergence time from closely related primate species.

## Results

### Estimation of mutation rate for 19 trios of rhesus macaques

To produce an accurate estimate for the germline mutation rate of rhesus macaques, we generated high coverage (76 X per individual after mapping, min 64 X, max 86 X) genome sequencing data for 19 trios of two unrelated families (Fig 1). The first family consisted of two reproductive males and four reproductive females, and the second family had one reproductive male and seven reproductive females. In the first family, the pedigree extended over a third generation in two cases. The promiscuous mating system of rhesus macaques allowed us to follow the mutation rates in various ages of reproduction, and compare numerous full siblings and half-siblings.

**Fig 1.**
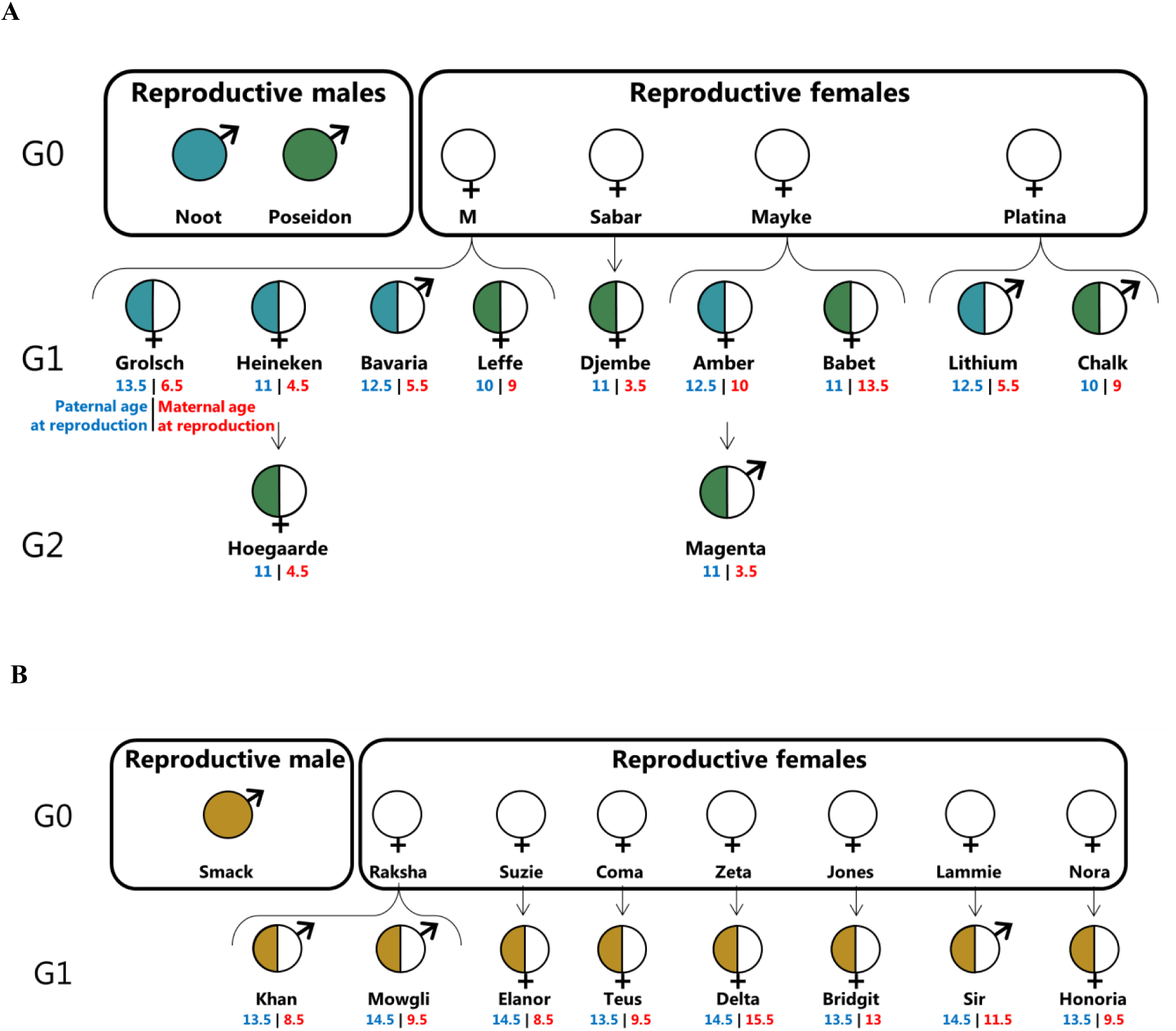
Pedigree of the 19 trios used for the direct estimation of mutation rate. A: The first group is composed of two reproductive males and four reproductive females. B: The second group contained one reproductive male and seven reproductive females. In each offspring, the color on the left corresponds to the paternal lineage and under the name are the age of the father (in blue) and mother (in red) at the time of reproduction. The reproductive ranges are 4.5 years for males and 12.2 years for females.

We developed a pipeline for single nucleotide polymorphisms (SNP) calling with multiple quality control steps involving the filtering of reads and sites (see Methods). For each trio, we considered candidate sites as *de novo* mutations when i) both parents were homozygotes for the reference allele, while the offspring was heterozygous with 30 % to 70 % of its reads supporting the alternative allele, and ii) the three individuals passed the depth and genotype quality filters (see Methods). These filters were calibrated to ensure a low rate of false-positives among the candidate *de novo* mutations.

We obtained an unfiltered set of 12,785,386 average candidate autosomal SNPs per trio (se = 26,196), of which a total of 177,227 were potential Mendelian violations (average of 9,328 per trio; se= 106). Of these, 744 SNPs passed the filters as *de novo* mutations, ranging from 25 to 59 for each trio and an average of 39 *de novo* mutations per trio (se = 2) (see S1 Table). We manually curated all mutations using IGV on bam files and found that 663 mutations convincingly displayed as true positives. This leaves a maximum of 10.89 % (81 sites) that could be false-positives due to the absence of the variant in the offspring or presence of the variant in the parents (see S1 Fig and the 81 curated mutations in supplementary). Most of those sites were in dinucleotide repeat regions or short tandem repeats (56 sites), while others were in non-repetitive regions of the genome (25 sites). The manual curation may have missed the realignment executed during variant calling. Thus, in the absence of objective filters, we decided to keep these regions in the estimate of mutation rate but corrected the number of mutations for each trio with a false-positive rate (see equation 1 in Methods section).

To confirm the authenticity of the *de novo* mutations, we performed PCR experiments for all candidate *de novo* mutations from one trio before manual correction. We designed primers to a set of 39 *de novo* candidates among which 3 *de novo* mutations assigned as spurious from the manual inspection. Of these, 24 sites were successfully amplified and sequenced for all three individuals i.e mother, father, and offspring, including 1 of the spurious sites. Among those sequenced sites, 23 were correct, only one was wrong (S2 Fig). This invalidated candidate was the spurious candidate removed by manual curation, therefore supporting our manual curation method. The PCR validation results suggested a lower false-positive rate of 4.2 % before manual curation. As the PCR validation was done only on 24 candidates we decided to keep a strict false-positive rate of 10.89 % found by manual curation.

We then estimated the mutation rate, per site per generation, as the number of mutations observed, and corrected for false-positive calls, divided by the number of callable sites. The number if callable sites for each trio ranged from 2,334,764,487 to 2,359,040,186, covering on average 88 % of the autosomal sites of the rhesus macaque genome. A site was defined as callable when both parents were homozygotes for the reference allele, and all individuals passed the depth and genotype quality filters at that site. As callability is determined using the base-pair resolution vcf file, containing every single site of the genome, all filters used during calling were taken into account during the estimation of callability, except for the site filters and the allelic balance filter. We then corrected for false-negative rates, calculated as the number of “good” sites that could be filtered away by both the site filters and allelic balance filters - estimated at 4.02 % (see equation 1 in Methods section). Another method to estimate the false-negative rate is to simulate mutations on the bam files and evaluate the detection rate after passing through all filters. On 552 randomly simulated mutations among the 19 offsprings, 545 were detected as *de novo* mutations, resulting in a false-negative rate of 1.27 %. The 7 remaining mutations were filtered away by the allelic balance filter only, which can be explained by the reads filtering in the variant calling step. This result might be underestimated due to the methodological limitation of simulating *de novo* mutations, yet, it ensures that a false-negative rate of 4.02 % is not out of range. Thus, the final estimated average mutation rate of the rhesus macaques was 0.77 × 10^−8^ *de novo* mutations per site per generation (95 % CI 0.69 × 10^−8^ - 0.85 × 10^−8^). We removed the 81 sites that, based on manual curation, could represent false-positive calls from the following analyses (see the 663 *de novo* mutations in S2 Table).

### Parental contribution and age impact to the *de novo* mutation rate

We observed a positive correlation between the paternal age and the mutation rate in the offspring (adjusted R^2^ = 0.23; *P* = 0.021; regression: *μ* = 1.022 × 10^−9^ + 5. 393 × 10^−10^ × *age*_*paternal*_; *P* = 0.021; Fig 2A). We also detected a slight positive correlation with the maternal age, though not significant (adjusted R^2^ = 0.09; *P* = 0.111; regression: *μ* = 6.200×10^−9^+1.818× 10^−10^×*age*_*maternal*_; P = 0.111; Fig 2B). A multiple regression of the mutation rate on paternal and maternal age resulted in this formula: *μ*_*Rhesus*_ = 1.355×10^−9^ +7.936×10^−11^ ×*age*_*maternal*_ +4.588× 10^−10^ × *age*_*paternal*_ (P = 0.06), where *μ*_*Rhesus*_ is the mutation rate for the species. We were able to phase 337 mutations to their parent of origin, which accounted for more than half of the total number of *de novo* mutations (663). There is a significant male bias in the contribution of *de novo* mutations, with an average of 80.6 % paternal *de novo* mutations (95 % CI 76.6 % - 84.6 %; T = 22.62, DF = 36, P < 2.2 × 10^−16^; Fig 2C). Moreover, with more than half of the *de novo* mutations phased to their parent of origin, we were able to disentangle the effect of the age of each parent on mutation rate independently (Fig 2D). By assuming that the ratio of mutations phased to a particular parent was the same in the phased mutations than in the unphased ones, we could predict the total number of mutations given by each parent. For instance, if an offspring had 40 *de novo* mutations and only half were phased, with 80 % given from its father, we would apply this ratio to the total number of mutations in this offspring, ending up with 32 *de novo* mutations from its father and eight from its mother. This analysis suggested a stronger male age effect to the number of mutations (adjusted R^2^ = 0.41, P = 0.002), and a similar, non significant maternal age effect (adjusted R^2^ = -0.01, P = 0.38). The two regression lines meet around the age of sexual maturity (3 years for females and 4 years for males), which is consistent with a similar accumulation of *de novo* mutations during the developmental process from birth to sexual maturity in both sexes, but the variances on the regression line slopes are large (see Fig 2C and S3 Fig for the same analysis with a Poisson regression). Using these two linear regressions, we can predict the number of *de novo* mutations in the offspring based on the age of each parent at the time of reproduction: *nb of mutations* _*Rhesus*_ = 4.6497+ 0.3042× *age*_*maternal*_ +4.8399 + 1.8364 × *age*_*paternal*_, where *nb of mutations* _*Rhesus*_ is the number of *de novo* mutations for the given trio. The expected mutation rates calculated using the two different regression models show similar correlations with the observed mutation rate (R^2^ = 0.54, P = 0.016 for the first regression and R^2^ = 0.54, P = 0.016 for the upscaled one, see S4 Fig). However, on the first regression on the mutation rate, the maternal age effect may be confounded by the paternal age, as maternal and paternal age are correlated in our dataset, yet, non-significantly (R^2^ = 0.38, P = 0.106, see S5 Fig). The upscaled regression unravels the effect of the parental age independently from each other. This regression can also be used to infer the contribution of each parent at different reproductive age. For instance, if both parents reproduce at 5 years old, based on the upscaled regression, the father is estimated to give ∽ 14 *de novo* mutations (95 % CI:6 - 22) and the mother ∽ 6 *de novo* mutations (95 % CI:3 - 10), corresponding to a contribution ratio from father to mother of 2.3:1 at 5 years old. If they reproduce at 15 years old, this ratio would be 3.6:1 with males giving ∽ 32 *de novo* mutations (95 % CI: 29 – 36) and females ∽ 9 *de novo* mutations (95 % CI: 4 – 14). It seems that the male bias increases with the parental age, yet, our model was based on too few data points in early male reproductive ages to reach a firm conclusion. For the two extended trios for which a second generation is available, we looked at the proportion of *de novo* mutations in the first offspring that were passed on to the third generation - the third generation inherited a heterozygote genotype with the alternative allele being the *de novo* mutation. In one case, 66 % of the *de novo* mutations in the female (Heineken) were passed to her daughter (Hoegaarde), while in another case, 40 % of the *de novo* mutations in the female (Amber) were passed to her son (Magenta). These deviations from the expected 50 % inheritance rate are not statistically significant (Binomial test; P_Hoegaarde_ = 0.14 and P_Magenta_ = 0.27).

**Fig 2.**
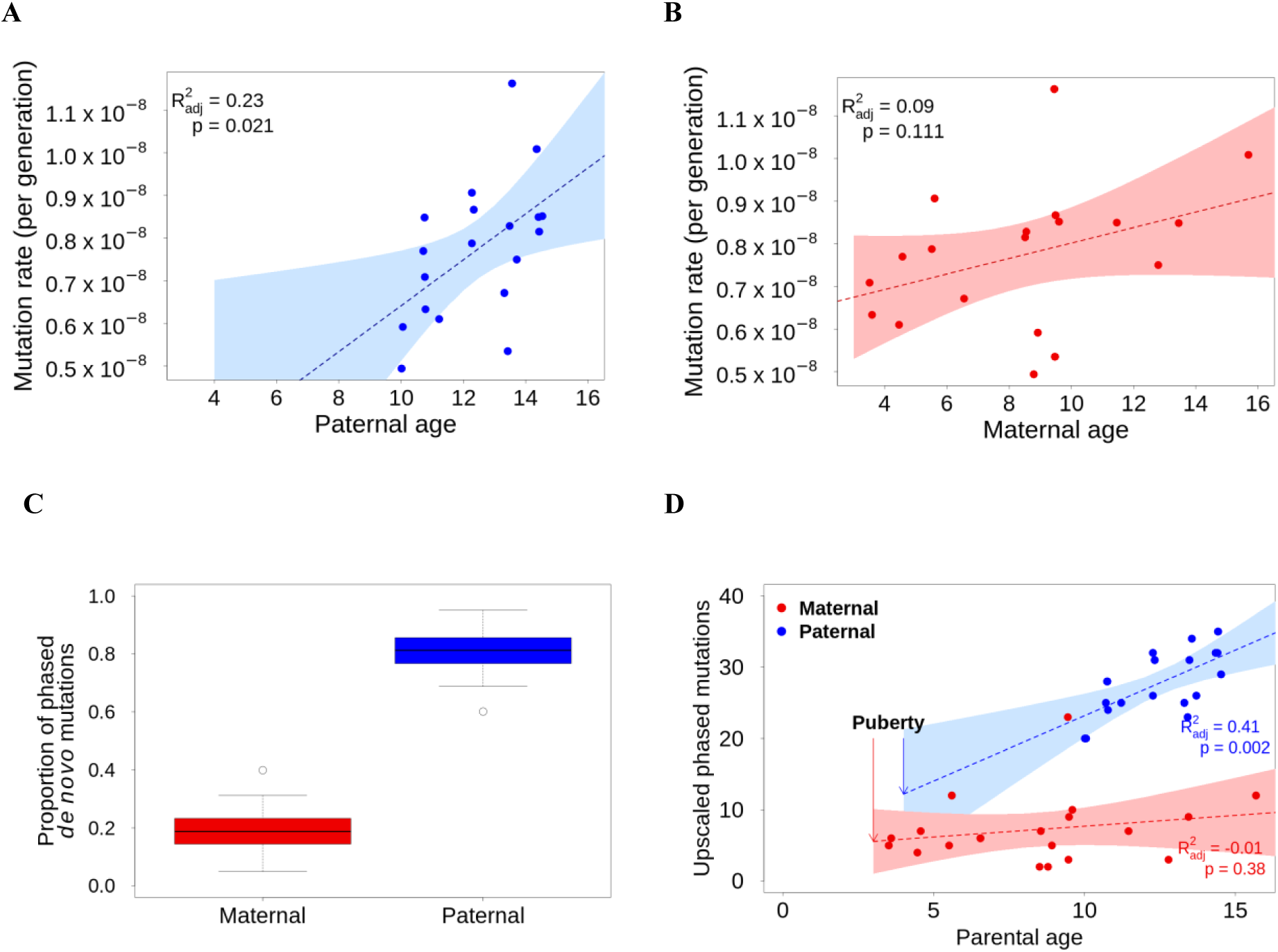
Parental contribution and age effect to the *de novo* mutation rate. A: There is a positive correlation between the mutation rate and the paternal age. B: The correlation between maternal age and mutation rate is not significant. C: Males contribute to 80.6 % of the *de novo* mutations while females contribute to 19.4 % of them. D: Upscaled number of *de novo* mutations given by each parent shows a similar contribution at the age of sexual maturation and a substantial increase with male age.

### Characterizations of *de novo* mutations

We characterized the type of *de novo* mutations and found that transition from a strong base to weak base (*G > A* and *C > T*) were most common (332/663), with 43 % of those mutations located in CpG sites (Fig 3A). In total, 23.2 % (154/663) of the *de novo* mutations were located in CpG sites. This is slightly higher than what has been found in humans, for which 19 % of the *de novo* mutations are in CpG sites (Besenbacher et al. 2015), but not significantly (human: X^2^ = 2.774, df = 1, P = 0.096). Moreover, 32.1 % (144/448) of the transition mutations (*A > G* and *C > T*) were in CpG sites, higher than what has been found in chimpanzee, with 24 % of the transition *de novo* mutations in CpG sites (Venn et al. 2014). The transition to transversion ratio (ti/tv) was 2.08, which is similar to the ratio observed in other species (human: ti/tv ∽ 2.16 (Yuen et al. 2016); human ti/tv ∽ 2.2 (Wang and Zhu 2014); chimpanzee: ti/tv ∽ 1.98 (Tatsumoto et al. 2017). The 663 *de novo* mutations showed some clustering in the genome (Fig 3B and S6 Fig). Across all trios, we observed 11 clusters, defined as windows of 20,000 bp where more than one mutation occurred in any individual, involving 23 mutations. Four clusters were made of mutations from a single individual, accounting for eight mutations (Fig 3B). Overall, 3.47 % of the *de novo* mutations were located in clusters, and 1.21 % were mutations within the same individual located in a cluster, which is significantly lower than the 3.1 % reported in humans (Besenbacher et al. 2016) (X^2^ = 7.35, DF = 1, P = 0.007; S7 Fig, S3 Table). We observed 23 mutations occurring recurrently in more than one related individual (Table 1), which accounted for 3.5 % of the total number of *de novo* mutations (23/663) and 1.5 % of sites (10/650 unique sites). Four *de novo* mutations (2 sites) were shared between half-siblings on the maternal side, and 19 (8 sites) were shared between half-siblings on the paternal side. However, there was no significant difference between the proportion of mutations shared between pairs of individuals related on the maternal side (9 pairs, 0.70 % shared), and pairs related on their paternal side (53 pairs, 0.80 % shared; Fisher’s exact test P = 1). In 6 sites, the phasing to the parent of origin confirmed that the mutation was coming from the common parent for at least one individual (Tab. 1). Moreover, the phasing was never inconsistent by attributing a shared *de novo* mutation to the other parent than the parent in common. However, 5 shared sites did appear as mosaic in the common parent, with a maximum of 5 % of the reads of the father supporting the alternative allele (4 out of 80 reads).. Nine of the *de novo* mutations (1.4 % of the total *de novo* mutations) were located in coding sequences (CDS regions), which is close to the overall proportion of coding sequences region (1.2%) in the whole macaque genome. Eigth out of those eight mutations were non-synonymous.

**Table 1.**
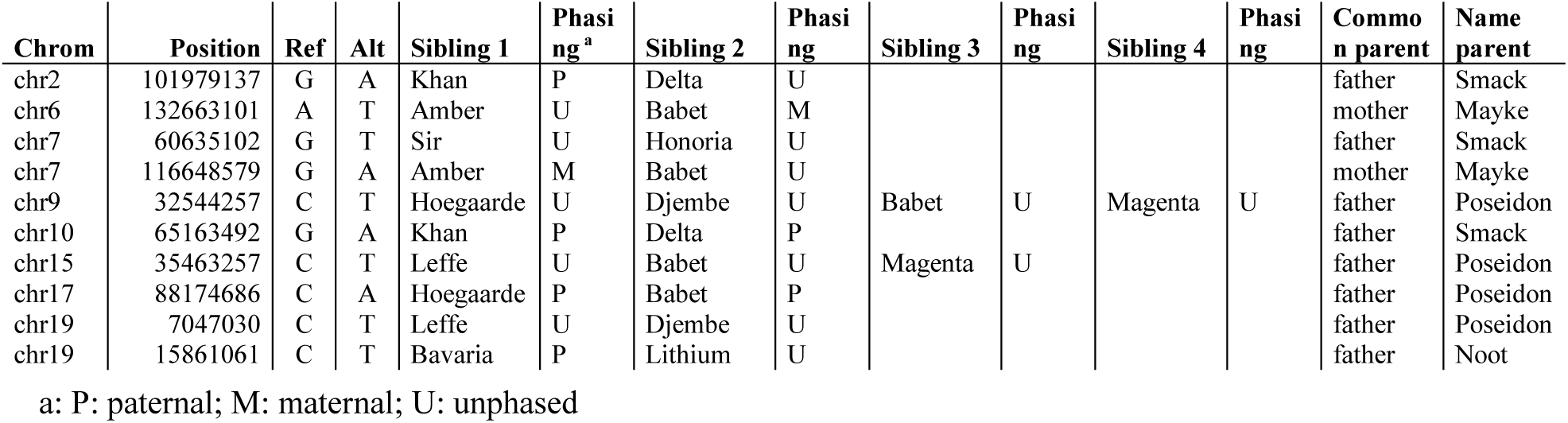
Six mutations shared between related individuals.

**Fig 3.**
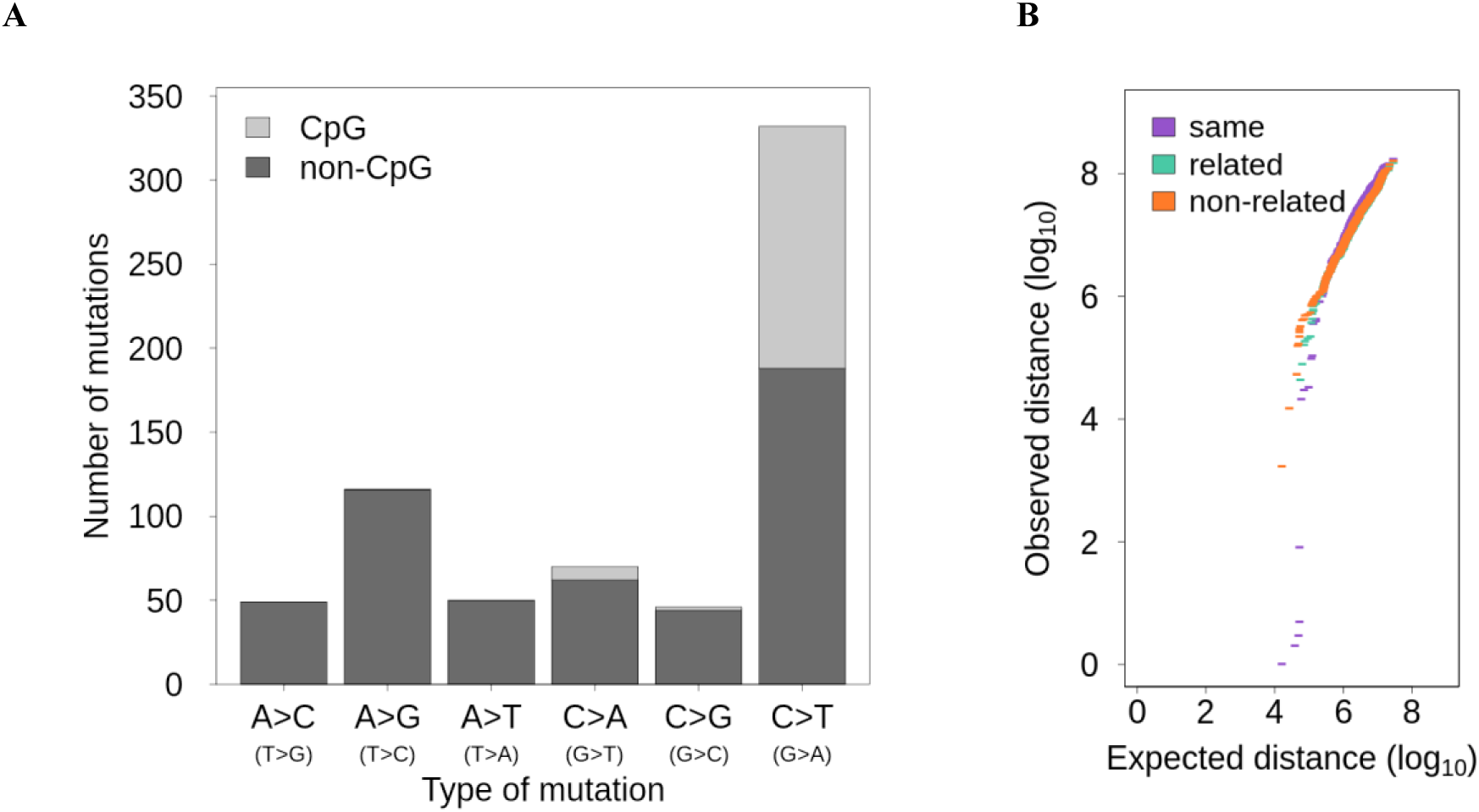
Characterizations of the *de novo* mutations. A: The type of *de novo* mutations in CpG and non-CpG sites. B: QQ-plot of the distance between *de novo* mutations compared to a uniform distribution within individuals (purple), between related individuals (green), and between non-related individuals (orange).

### Molecular dating with trio-based mutation rate

Based on our inferred mutation rate and the genetic diversity of Indian rhesus macaques (*π* = 0.00247) estimated using whole genomic sequencing data from more than 120 unrelated wild individuals (Xue et al. 2016), we calculated the effective population size (N_e_) of rhesus macaques to be 79,874. This is similar to the N_e_ = 80,000 estimated previously using *μ =* 0.59 × 10^−8^ from hippocampal transcriptome and H3K4me3-marked DNA regions from 14 individuals (Yuan et al. 2012), yet higher than N_e_ = 61,800 estimated using *μ =* 1×10^−8^ with 120 individual full genome data (Xue et al. 2016). Assuming a generation time of 11 years and an average reproduction age of 10 years for females and 12 years for males, the yearly mutation rate of rhesus macaques was calculated based on our regression model of the number of mutations given by males and females independently, and the average callability (see equation 2 in the Methods section). As captive animals usually reproduce later than in the wild, which could impact the average mutation rate per generation, we used the regression instead of the mutation rate per generation to correct for this possible bias. The yearly mutation rate of rhesus macaques in our calculation was 0.62 × 10^−9^ per site per year, almost 1.5 times that of humans (Jónsson et al. 2017).

Given a precise evolutionary mutation rate is essential for accurate calibration of molecular divergence events between species, we used the mutation rate we inferred for rhesus macaques to re-date the phylogeny of closely-related primate species with full genome alignment available (Moorjani et al. 2016) (Fig 4A). The molecular divergence time (T_D_) is the time since an ancestral lineage started to split into two descendant lineages, and can be inferred from the genetic divergence between the two descendant lineages and the mutation rate. The speciation time (T_S_) is a younger event that implies no more gene flow between lineages (Steiper and Young 2008). On the backward direction, the alleles of two descendant lineages are randomly sampled from their parents until going back to the most recent common ancestor (Rosenberg and Nordborg 2002). This stochastic event, known as the coalescent, depends on the population sizes, being slower in a large population (Kingman, 1982). Thus, from the divergence time, the speciation time can be inferred given the rate of coalescence (see equation 3 in the Method section). We also compared our results to those of previous dating attempts based on molecular phylogenetic trees calibrated with fossils records (Fig 4B). We found that the two methods concur for the most recent events. Specifically, we estimated that the *Macaca mulatta* and *Macaca fascicularis* genomes had already diverged around 3.90 million years ago (Mya) (95 % CI: 3.46 – 4.46), which is slightly older than previous estimates using the molecular clock calibrated with fossils, as the molecular divergence of the two species has been estimated at 3.44 Mya with mitochondrial data (Pozzi et al. 2014) and 3.53 Mya from nuclear data (Perelman et al. 2011).

We estimated a speciation event between the two species 2.14 Mya after the coalescent time, also consistent with previous findings of a most common recent ancestor to the two populations of the rhesus macaque, the Chinese and the Indian population, around 1.94 Mya based on coalescent simulations (Hernandez et al. 2007). For the next node, the molecular clock seems to differ between mitochondrial and nuclear data, as the divergence time for the Papionini group into the *Papio* and *Macaca* genera has been estimated to 8.13 Mya using nuclear data (Perelman et al. 2011), and 12.17 Mya with mitochondrial data (Pozzi et al. 2014). We estimated a divergence time between these two genera of 13.17 Mya (95 % CI: 11.70 – 15.07). For earlier divergence events, our estimated divergence times are more ancient than previous reports. For instance, we estimated that the Cercopithecini and Papionini diverged 19.86 Mya (95 % CI: 17.64 – 22.71), while other studies had calculated 11.55 Mya using nuclear data (Perelman et al. 2011), and 14.09 Mya using mitochondrial data (Pozzi et al. 2014). Finally, the divergence between Cercopithecidae and Hominoidea has been reported between 25 and 30 Mya (Stewart and Disotell 1998; Moorjani et al. 2016), with an estimation of 31.6 Mya using the nuclear molecular clock (Perelman et al. 2011) and 32.12 Mya using the mitochondrial one (Pozzi et al. 2014). Our dating of the divergence time between the Cercopithecidae and Hominoidea of 52.31 Mya (95 % CI: 46.47 – 59.83) is substantially older than previous estimates. However, the estimated speciation time inferred based on the ancestral population size, suggested a speciation of the Catarrhini group into two lineages 44.50 Mya (Fig 4B).

**Fig 4.**
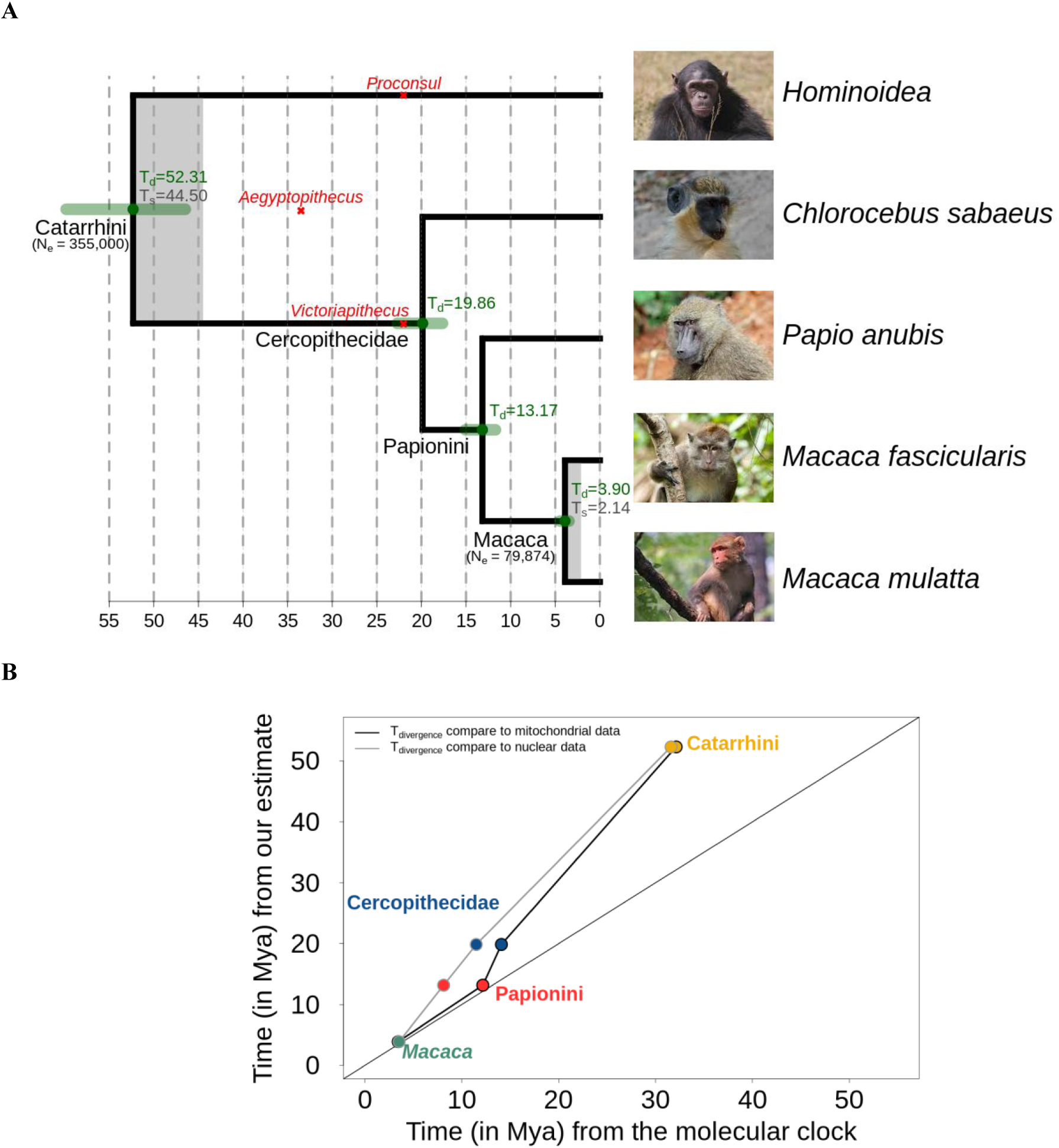
Molecular dating with pedigree-based mutation rate. A: Primates phylogeny based on the yearly mutation rate (0.62 × 10^*−*9^ per site per year). In green are the confidence interval of our divergence time estimates (Td) and grey shades represent the time of speciation (Ts). The effective population sizes are indicated under the nodes (N_e_ Macaca ancestor is our estimate of N_e_ *Macaca mulatta* and N_e_ Catarrhini from the literature (Schrago 2014)). B: Comparison of our divergence time and speciation time with the previous estimation using the molecular clock from mitochondrial (Pozzi et al. 2014) and nuclear data (Perelman et al. 2011) calibrated with fossils records.

## Discussion

Despite many efforts to accurately estimate direct *de novo* mutation rates, it is still a challenging task due to the rare occurrence of *de novo* mutations, and the small sample size that is often available. Sequencing coverage is known to be a significant factor in affecting false-positive (FP), and false-negative (FN) calls when detecting *de novo* mutation (Acuna-Hidalgo et al. 2016; Tatsumoto et al. 2017). A minimal sequencing coverage at 15X was recommended for SNPs calling (Song et al. 2016). However, such coverage cannot provide sufficient power to reduce FPs because the lower depth threshold cannot preclude Mendelian violations due to sequencing errors. Moreover, a larger portion of the genome would be removed in the denominator at low depth in order to reduce the FN. While most studies on direct estimation of mutation rate use 35-40X coverage (Jónsson et al. 2017; Thomas et al. 2018; Besenbacher et al. 2019), their methods to reduce FP and FN differ. Some studies use the deviation from 50 % of the *de novo* mutation pass to the next generation to infer the false-positive rate (Jónsson et al. 2017; Thomas et al. 2018). Others use probabilistic methods to access the callability (Besenbacher et al. 2019), or simulation of known mutation to control the pipeline quality (Pfeifer 2017). Differences in methods likely impact the calculated rate. Here, we produced sequences at 76X coverage, which allows us to apply conservative filtering processes, while still obtaining high coverage (88 %) of the autosomal genome region when inferring *de novo* mutations. To our knowledge, only one other study has used very high coverage (120X per individuals), on a single trio of chimpanzees (Tatsumoto et al. 2017). Such high coverage allowed us to achieve a false-positive rate below 10.89 % and within the regions we deemed callable, we calculated a low false-negative rate of 4.02 %.

Our estimated rate is higher than the 0.58 × 10^−8^ *de novo* mutations per site per generation estimated in a preprint report (Wang et al. 2020). The difference should be mainly attributed to the fact that they sequenced the offspring of younger parents (average parental age of 7.1 years for females and 7.8 years for males compared to 8.4 years for females and 12.4 years for males in this study). Using our regression from the phased mutation, we estimated a mutation rate of 0.51 × 10^−8^ per site per generation, when males reproduce at 7.8 years and females reproduce at 7.1 years old. Moreover, using their regression based on the age of puberty and the increase of paternal mutation per year, Wang and collaborators estimated a per generation rate of 0.71 × 10^−8^ mutations when males reproduce at 11 years, and a yearly rate of 0.65 × 10^−9^ mutations per site per year, which is approx 5 % higher than our estimate of 0.62 × 10^−9^ (2020). This difference may be due to any combination of stochasticity, differences in *de novo* mutation rate pipelines (callability estimate, false-negative rate, and false-positive rate estimate) and different models for converting pedigree estimates to yearly rates. Our combination of high coverage data and a large number of trios allowed us to gain high confidence estimates of the germline mutation rate of rhesus macaques at around 0.77 × 10^−8^ *de novo* mutation per site per generation, ranging from 0.49 × 10^−8^ to 1.16 × 10^−8^. This is similar to the mutation rate estimated for other non-Hominidae primates; 0.81 × 10^−8^ for the owl monkey (*Aotus nancymaae*) (Thomas et al. 2018) and 0.94 × 10^−8^ for the African green monkey (*Chlorocebus sabaeus*) (Pfeifer 2017), while all Hominidae seem to have a mutation rate that is higher than 1 × 10^−8^ *de novo* mutation per site per generation (Jónsson et al. 2017; Besenbacher et al. 2019). However, if we count for the *de novo* mutation per site per year, the rate of rhesus macaque (0.62 × 10^−9^) is almost 1.5-fold the human one of 0.43 × 10^−9^ mutation per sites per year (Jónsson et al. 2017).

One of the main factors affecting the mutation rate within the species is the paternal age at the time of reproduction, which was attributed to the accumulation of replication-driven mutations during spermatogenesis (Drost and Lee 1995; Li et al. 1996; Crow 2000), and has been observed in many other primates (Venn et al. 2014; Jónsson et al. 2017; Maretty et al. 2017; Thomas et al. 2018; Besenbacher et al. 2019). In rhesus macaques, the rate at which germline mutation increases with paternal age seems faster than in humans; we inferred 1.84 mutations more per year for the rhesus macaque father (95% CI 0.77 – 2.90 for an average callable genome of 2.35 Mb), compared to 1.51 in humans (95% CI 1.45–1.57 for an average callable genome of 2.72 Mb) (Jónsson et al. 2017). For females, there is less difference, with 0.30 more mutations per year for the mother in rhesus macaque (95% CI -0.41 – 1.02), and 0.37 more per year in human mothers (95% CI 0.32–0.43) (Jónsson et al. 2017). In rhesus macaques, males produce a larger number of sperm cells per unit of time (23 × 10^6^ sperm cells per gram of testis per day (Amann et al. 1976)) than humans (4.4 × 10^6^ sperm cells per gram of testis per day (Amann and Howards 1980)). This could imply a higher number of cell division per unit of time in rhesus macaques and thus more replication error during spermatogenesis. This is also consistent with the generation time effect which stipulates that an increase in generation time would decrease the number of cell division per unit of time as well as the yearly mutation rate assuming that most mutations arise from replication errors (Wu and Lit 1985; Goodman et al. 1993; Ohta 1993; Li et al. 1996; Ségurel et al. 2014; Scally 2016). Indeed, humans have a generation time of 29 years, while it is 11 years for rhesus macaques. Another explanation for a higher increase of mutation rate with paternal age could be differences in the replication machinery itself. Due to higher sperm competition in rhesus macaque, the replication might be under selective pressure for fast production at the expense of replication fidelity, leading to less DNA repair mechanisms. As in other primates, we found a male bias in the contribution of *de novo* mutations, as the paternal to maternal ratio is 4.2:1.This ratio is higher than the 2.7:1 ratio observed in mice (Lindsay et al. 2019) and slightly higher than the 4:1 ratio observed in humans (Goldmann et al. 2016; Jónsson et al. 2018; Lindsay et al. 2019). Similarly to the wild, the males of our dataset reproduced from 10 years old, which did not allow us to examine if the contribution bias was also present just after maturation. Moreover, the promiscuous behavior of rhesus macaque leads to father reproducing with younger females. Using our model to compare the contribution of each parent reproducing at the similar age, it seems that the male bias increases with the parental age, with a lower difference in contribution at the time of sexual maturation (2.3:1 for parents of 5 years old) and an increase in male to female contribution with older parents (3.6:1 for parents of 15 years old). This result differs from humans, where the male bias seems constant over time (Gao et al. 2019), but more time points in macaque would be needed to interpret the contribution over time. In rhesus macaques, the ratio of paternal to maternal contribution to the shared mutations between related individuals is 1:1, similarly to what has been shown in mice (Lindsay et al. 2019), highlighting that those mutations probably occur during primordial germ cell divisions in postzygotic stages. Our study shows many shared patterns in the *de novo* mutations among non-Hominid primates. More estimation of mammals could help understanding if these features are conserved across a broad phylogenetic scale. Moreover, further work would be needed to understand if some gamete production stages are more mutagenic in some species than others.

An accurate estimation of the mutation rate is essential for the precise dating of species divergence events. We used the rhesus macaque mutation rate to estimate its divergence time with related species for which whole-genome alignments are already available and their molecular divergence times have been investigated before with other methods (Moorjani et al. 2016). The results of our direct dating method, based on molecular distances between species and *de novo* mutation rate, matched those of traditional molecular clock approaches for speciation events within 10 to 15 million years. However, it often produced earlier divergence times for more ancient nodes than the molecular clock method. This incongruence might be attributed to the fossils that were used for calibration with the clock method, which has many limitations (Heads 2005; Pulquério and Nichols 2007; Steiper and Young 2008). A fossil used for calibrating a node is usually selected to represent the oldest known specimen of a lineage. Still, it cannot be known if real even older specimens existed (Heads 2005). Thus, a fossil is usually assumed to be younger than the real divergence time of the species (Benton et al. 2015). Moreover, despite the error associated with the dating of a fossil itself, determining its position on a tree can be challenging and have effects on the inferred ages across the whole tree (Pulquério and Nichols 2007; Steiper and Young 2008). For instance, the Catarrhini node, marking the divergence between the Cercopithecidae and the Hominoidea, is often calibrated in primate phylogenies (Heads 2005). This node has been calibrated to approx. 25 Mya using the oldest known Cercopithecidae fossil (*Victoriapithecus*), and the oldest known Hominoidea fossil (*Proconsul*), both around 22 My old (Goodman et al. 1998). However, if the oldest Catarrhini fossil (*Aegyptopithecus*) of 33 to 34 My age is used, this node could also be calibrated to 35 Mya (Stewart and Disotell 1998). Finally, instead of being an ancestral specimen of the Catarrhini, *Aegyptopithecus* has been suggested as a sister taxon to Catarrhini, which would lead to an even older calibration time for this node (Stewart and Disotell 1998).

On the other hand, the direct mutation rate estimation could have produced overestimated divergence times for the Catarrhini node age compared to previous estimates (Perelman et al. 2011; Pozzi et al. 2014), because the mutation rate and generation time might change cross-species and over time. It is possible that the Catarrhini ancestor would have had a faster yearly mutation rate, and/or a shorter generation time than the recent macaques. Since fossil calibration could underestimate real divergence times, molecular-based methods could overestimate it, especially by assuming a unique mutation rate to an entire clade (Steiper and Young 2008).

To obtain more confidence in the estimation of divergence time, it would be necessary to have an accurate estimation of the mutation rate for various species. The estimates available today for primates vary from 0.81×10^−8^ per site per generation for the Owl monkey (*Aotus nancymaae*) to 1.66×10^−8^ per site per generation for Orangutan (*Pongo abelii*). However, the different methods and sequencing depth make it difficult to compare between species and attribute differences to biological causes or methodological ones. Therefore, more standardized methods in further studies would be needed to allow for cross-species comparison.

## Methods

### Samples

Whole blood samples (2 mL) in EDTA (Ethylenediaminetetraacetic acid) were collected from 53 Indian rhesus macaques (Macaca mulatta) during routine health checks at the Biomedical Primate Research Centre (BPRC, Rijswijk, Netherlands). Individuals originated from two groups, with one or two reproductive males per group. After ensuring the relatedness with a test based on individual genotypes (Manichaikul et al. 2010), we ended up with 19 trios formed by 33 individuals and two extended trios (for which a second generation was available). In our dataset males reproduced from 10 years old to 14.5 years old (♂ reproductive range: 4.5 years), and females from 3.5 years old to 15.7 years old (♀ reproductive range: 12.2 years). Genomic DNA was extracted using DNeasy Blood and Tissue Kit (Qiagen, Valencia-CA, USA) following the manufacturer’s instructions. BGIseq libraries were built in China National GeneBank (CNGB), Shenzhen, China. The average insert size of the samples was 230 base pairs. Whole-genome pair-ended sequencing was performed on BGISEQ500 platform, with a read length of 2×100 bp. The average coverage of the raw sequences before trimming was 81X per sample (SE = 1.35). Whole-genome sequences have been deposited in NCBI (National Center for Biotechnology Information) with BioProject number PRJNA588178 and SRA submission SUB6522592.

### Reads mapping, SNPs calling, and filtering pipeline

Adaptors, low-quality reads, and N-reads were removed with SOAPnuke filter (Chen et al. 2017). Trimmed reads were mapped to the reference genome of rhesus macaque Mmul 8.0.1 using BWA-MEM version 0.7.15 with the estimated insert size option. Only reads mapping uniquely were kept and duplicates were removed using Picard MarkDuplicates. The average coverage after mapping was 76X per individuals (SE = 1.16). Variants were called using GATK (Poplin et al. 2018); calling variants for each individual with HaplotypeCaller in BP-RESOLUTION mode; all gVCF files per sample were combined into a single one per trio using CombineGVCFs per autosomal chromosomes; finally joint genotyping was applied with GenotypeGVCF. Because *de novo* mutations are rare events, variant quality score recalibration (VQSR) is not a suitable tool to filter the sites as *de novo* mutations are more likely to be filtered out as low-quality variants. Instead we used a site filtering with the following parameters: QD < 2.0, FS > 20.0, MQ < 40.0, MQRankSum < - 2.0, MQRankSum > 4.0, ReadPosRankSum < - 3.0, ReadPosRankSum > 3.0 and SOR > 3.0. These filters were chosen by first, running the pipeline with the site filters recommended by GATK (QD < 2.0; FS > 60.0; MQ < 40.0; MQRankSum < -12.5; ReadPosRankSum < -8.0; SOR > 3.0), then, doing a manual curation of the candidates *de novo* mutations on the Integrative Genome Viewer (IGV). Finally, we identified the common parameters within the apparent false-positive calls and decided to adjust the site filter to remove as many false-positives without losing much true positive calls (see the pipeline S8 Fig).

### Detection of *de novo* mutations

The combination of high coverage (76X) and stringent filters reduced false-positive - calling a *de novo* mutation while it is not there. Thus, for each trio, we applied the following filters:

a. Mendelian violations were selected using GATK SelectVariant and refined to only keep sites where both parents were homozygote reference (HomRef), and their offspring was heterozygote (Het).
b. In the case of a *de novo* mutation, the number of alternative alleles seen in the offspring should account for ∽ 50 % of the reads. Our allelic balance filter allowed the alternative allele to be present in 30 % to 70 % of the total number of reads (applying the same 30% cutoff as in other studies (Kong et al. 2012; Besenbacher et al. 2015; Francioli et al. 2015; S9 Fig).
c. The depth of the three individuals was filtered to be between 0.5×*m*_*depth*_ and 2×*m*_*depth*_, with *m*_*depth*_ being the average depth of the trio. Most of the Mendelian violations are due to sequencing errors in regions of low sequencing depth; therefore, we applied a stricter threshold on the minimum depth to avoid the peak of Mendelian violations around 20X (S10 Fig).
d. Finally, after analyzing each trio with different genotype quality GQ cutoff (from 10 to 90), we set up a filter on the genotype quality of 60 to ensure the genotypes of the HomRef parents and the Het offspring (S11 Fig).

From 242,922,329 autosomal SNPs (average of 12,785,386 per trio), 2,251,363 were potential Mendelian violations found by GATK (average of 118,493 per trio), 177,227 were filtered Mendelian violations with parents HomRef and offspring Het (average of 9,328 per trio) (a), 78,339 passed the allelic balance filter (average of 4,123 per trio) (b), 13,251 passed the depth filter (average of 697 per trio) (c) and 744 the genotype quality filter (average of 39 per trio) (d) (see S4 Table for details on each individual). We also remove sites where a *de novo* mutation was shared among non-related individuals (1 site shared between 4 unrelated individuals). This allowed us to detect the number of *de novo* mutations observed per trio called m. We manually checked the reads mapping quality for all *de novo* mutations sites in the Integrative Genome Viewer (IGV). And we found possible false-positive calls in 10.89 % of the sites for which the variant was absent from the offspring or also present in a parent (see S1 Fig). We kept those sites for the estimation of the mutation rate, and corrected for false-positive (*β* = 0.1089), but removed them for downstream pattern analysis. We experimentally validated the *de novo* candidates from the trio Noot (father), Platina (mother), and Lithium (offspring). Primers were designed for 39 candidates (S5 Table). PCR amplification and Sanger sequencing were conducted on each individual (protocol in Supplementary materials). On 24 sites the PCR amplification and sequencing returned high-quality results for all three individuals. A candidate was considered validated when both parents showed homozygosity for the reference allele and the offspring showed heterozygosity (S2 Fig). All sequences generated for the PCR validation have been deposited in Genbank with accession numbers MT426016 - MT426087 (S4 Table).

### Estimation of the mutation rate per site per generation

From the number of *de novo* mutations to an estimate of the mutation rate per site per generation, it is necessary to also correct for false-negatives - not calling a true *de novo* mutation as such. To do so, we estimated two parameters: the false-negative rate and the number of callable sites, *C*, ie. the number of sites in the genome where we would be able to call a *de novo* mutation if it was there. We used the BP_RESOLUTION option in GATK to call variants for each position and thus get the exact genotype quality for each site in each individual - also sites that are not polymorphic. So unlike other studies, we do not have to rely on sequencing depth as a proxy for genotype quality at those sites. Instead, we can apply the same genotype quality threshold to the non-polymorphic sites as we do for *de novo* mutation candidate sites. This should lead to a more accurate estimate of the number of callable sites. For each trio, *C* is the sum of all sites where: both parents are HomRef, and the three individuals passed the depth filter (b) and the genotype quality filter (d). To correct for our last filter, the allelic balance (c), we estimated the false-negative rate *α*, defined as the proportion of true heterozygotes sites (one parent HomRef, the other parent HomAlt and their offspring Het) outside the allelic balance threshold (S9 Fig). We also implemented in this parameter the false-negative rate of the site filters following a normal distribution (FS, MQRankSum, and ReadPosRankSum). For all trios combined, the rate of false-negatives caused by the allele balance filter and the site filters was 0.0402. To validate this false-negative rate estimation we also used a simulation method, used in other studies (Keightley et al. 2015; Pfeifer 2017). With BAMSurgeon (Ewing et al. 2015), 552 mutations were simulated across the 19 trios at random callable sites. The false-negative rate was calculated as 1 – (number of detected mutations/number of simulated mutations), after running the pipeline from variant calling. The mutation rate per sites per generation can then be estimated per trio with the following equation:

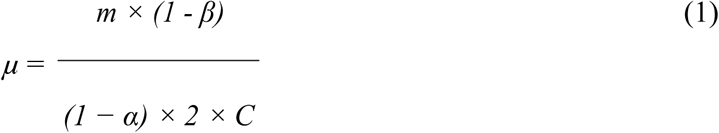

### Sex bias, ages, and relatedness

*De novo* mutations were phased to their parental origin using the read-backed phasing method described in Maretty et al. 2017 (script available on GitHub: https://github.com/besenbacher/POOHA). The method uses read-pairs that contain both a *de novo* mutation and another heterozygous variant, the latter of which was used to determine the parental origin of the mutation if it is present in both offspring and one of the parents. The phasing allowed us to identify any parental bias in the contribution of the *de novo* mutations. Pearson’s correlation test was performed between the mutation rate and ages of each parent, as well as a linear regression model for father and mother independently. A multiple linear regression model was performed to predict the mutation rate from both parental ages as predictor variables. The phased mutations were used to dissociate the effect of the parental age from one another. Because the total number of SNPs phased to the mother or the father may differ, we divided the phased *de novo* mutations found in a parent by the total SNPs phased to this parent. Only a subset of the *de novo* mutations in an offspring was phased. Thus, we applied the paternal to maternal ratio to the total number of mutations in a trio, referred to as ’upscaled’ number of mutations, to predict the number of total mutations given by each parent at different ages. The two extended trios, analyzed as independent trios, also allowed us to determine if ∽ 50 % of the *de novo* mutations observed in the first trio were passed on to the next generation.

### Characterization of *de novo* mutations

From all the *de novo* mutations found, the type of mutations and their frequencies were estimated. For the mutations from a C to any base we determined if they were followed by a G to detect the CpG sites (similarly if G mutations were preceded by a C. We defined a cluster as a window of 20,000 bp to qualify how many mutations were clustered together; over all individuals, looking at related individuals, and within individuals. We simulated 663 mutations following a uniform distribution to compare with our dataset. We investigated the mutations that are shared between related individuals. Finally, we looked at the location of mutations in the coding region using the annotation of the reference genome.

### Molecular dating using the new mutation rate

We calculated the effective population size using Watterson’s estimator *θ* = 4*N*_*e*_*μ* (Watterson 1975). We estimated *θ* with the nucleotide diversity *π* = 0.00247 according to a recent population study (Xue et al. 2016). Thus, we calculated the effective population size as 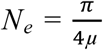 with *μ* the mutation rate per site per generation estimated in our study. To calculate divergence time, we converted the mutation rate to a yearly rate based on the regression model of the number of mutations given by each parent regarding their ages and the average callability C = 2,351,302,179. Given the maturation time and the high mortality due to predation, we assumed an average age of reproduction in the wild at 10 years old for females and 12 years old for males and a generation time of 11 years, also reported in another study (Xue et al. 2016). Thus, the yearly mutation rate was:

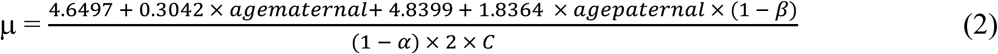

The divergence time between species was then calculated using 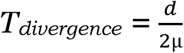 with *d* the genetic distance between species which were calculated from the whole-genome comparison (Moorjani et al. 2016) and *μ* the yearly mutation rate of rhesus macaques. We also used the confidence interval at 95% of our mutation rate regression to compute the confidence interval on divergence time. Based on the coalescent theory (Kingman, 1982), the time to coalescence is 2NeG with G the generation time and Ne the ancestral effective population size, assumed constant over time, as shown in a previous study (Xue et al. 2016). Thus, we dated the speciation event as previously done by Besenbacher et al. 2019 with:

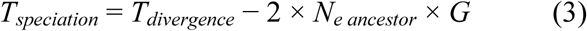

## Acknowledgments

We would like to thank GenomeDK at Aarhus University for providing computational resources and supports to this study. We also thank Josefin Stiller for helpful comments on the manuscript.

## Data Availability Statement

Whole-genome sequences have been deposited in NCBI (National Center for Biotechnology Information) with BioProject number PRJNA588178 and SRA submission SUB6522592. All sequences generated for the PCR validation have been deposited in Genbank with accession numbers MT426016 - MT426087.

